# Hydroxychloroquine alters the cytoskeleton to impair cell migration

**DOI:** 10.64898/2026.05.16.725640

**Authors:** Dipanjan Ray, Asim Bisoi, Bipasa Chaudhuri, Prashant Chandra Singh, Deepak Kumar Sinha

## Abstract

Hydroxychloroquine (HCQ), a clinically relevant quinoline derivative, impairs both collective and individual cell migration by disrupting actin and vimentin cytoskeletal dynamics during wound healing. In scratch assays with HeLa cells, HCQ treatment significantly reduces wound closure rates and inhibits bursts of coordinated migration, as well as single-cell motility. Quantitative imaging reveals that HCQ diminishes actin filament density at the wound edge and induces reorganization of the vimentin network, resulting in smaller nuclei and compromised structural connectivity. Particle tracking micro-rheology demonstrates that HCQ softens the cytoplasm and decreases cellular mechanical heterogeneity. *In vitro* spectroscopic studies show that HCQ binds cooperatively to actin with micromolar affinity, perturbing its secondary structure, reducing filament polymerisation rates, and impairs interactions between actin and actin binding proteins (ABPs). HCQ also significantly suppresses lamellipodial protrusive activity, indicating a link between cytoskeletal remodelling and impaired cell migration. Collectively, these findings establish that HCQ disrupts essential mechanisms for directed migration by modulating the cytoskeleton and cell mechanics. This multifaceted impairment may underlie therapeutic potential of HCQ as an anticancer agent by restricting cellular invasive capacity and remodelling required for tumour progression.

## Introduction

Cell migration under physiological conditions is a dynamic and highly coordinated biological process that allows cells to relocate to distinct spatial positions in response to diverse internal and external stimuli. The migration is associated with several biological processes such as cell growth and differentiation ^1^, morphogenesis ^2^, signal transduction ^3^, tissue organization ^4^, immune surveillance ^5^ and many more. At the molecular level, cell migration follows cyclical sequences of events such as polarization, protrusion formation, adhesion, contraction, and detachment ^6^. All these steps are closely associated with cytoskeletal remodelling, which provides the structural framework and generate the mechanical forces required for cell motility. Cytoskeletal reorganization during cell migration in physiological condition is highly coordinated and transient, ensuring control over cell shape, polarity, and movement. Actin filaments, microtubules, and intermediate filaments remodel in response to regulated signalling cues, allowing cells to migrate only when and where needed, such as during tissue repair or immune surveillance. This remodelling is reversible and balanced, maintaining overall tissue structure and homeostasis ^7,8^.

In contrast to physiological condition, the cell migration becomes uncontrolled and dysregulated in cancerous condition ^9^. Cancer cells often acquire abnormal motility through factors such as changes in signalling pathways, loss of cell-cell adhesion, and reorganization of the cytoskeleton ^10,11^. In cancer conditions, however, cytoskeletal reorganization becomes aberrant and persistent due to the activation of oncogenic signalling pathways ^10,11^. These alterations lead to excessive actin polymerization ^12^, increased focal adhesion turnover ^13^, and enhanced actomyosin contractility ^14^, which collectively promote uncontrolled motility and invasiveness. As a result, cancer cells exhibit abnormal morphology ^12^, lose polarity ^15^, and acquire migratory phenotypes such as epithelial-mesenchymal transition (EMT), enabling metastasis ^14^. Hence, one of the primary objectives of cancer drugs is to modulate cell migration through various mechanisms ^16^, as suppressing the migratory and invasive behaviour of tumour cells is critical for preventing metastasis ^17^, which is the leading cause of cancer-related mortality. Hydroxychloroquine (HCQ) is a well-known immunomodulatory agent widely used to treat rheumatoid arthritis, systemic lupus erythematosus, and malaria ^18–20^. Recent studies have demonstrated that HCQ also exerts significant effects on cancer cells, including enhancing tumour sensitivity to existing therapies ^21,22^. Consequently, numerous clinical trials have been initiated to evaluate HCQ as a novel targeted agent against various cancer types, with over thirty studies currently registered across all phases of clinical evaluation ^23^. Importantly, HCQ has been shown to have a stronger impact on gene regulation in cancer cell lines compared to chloroquine and other analogues ^24,25^, highlighting its potential as a promising candidate for cancer therapy, either as a stand-alone treatment or in combination with other drugs. Given these effects, it is plausible that HCQ may also influence the migration of cancer cells, thereby potentially limiting disease progression and metastasis.

In this study, we aimed to elucidate the impact of HCQ on the migratory behaviour of cancer cells, focusing specifically on the underlying cytoskeletal remodelling processes using array of in vitro and in vivo biophysical and biochemical techniques. Our data suggests that HCQ markedly impairs both collective and single-cell migration by disrupting actin-driven protrusive activity. HCQ reduces wound-closure rates, suppresses lamellipodial dynamics, diminishes cell motility, and alters directional persistence. It reorganizes cytoskeletal architecture by decreasing actin filament density and inducing vimentin redistribution, leading to cytoplasmic softening. In-vitro study reveals that HCQ binds cooperatively to actin with micromolar affinity, perturbs its secondary structure, slows polymerization, and weakens interactions with actin-binding proteins (ABPs) such as α-actinin. Together, these effects compromise cytoskeletal integrity, ultimately restricting cell movement.

## Results

### HCQ impairs collective cell migration during wound healing

Given the growing clinical interest in quinoline based molecules as potential anticancer agents, we examined the effects of HCQ, an FDA approved drug, on cell migration during the wound healing. To determine whether hydroxychloroquine (HCQ) induces cytotoxicity during short-term treatment, HeLa cells were treated with 100 µM HCQ for the indicated durations (0, 8, 16, and 24 h), and cell viability was measured using the MTT assay (**Figure S1-A**). HCQ treatment resulted in a gradual time-dependent decrease in MTT reduction. Importantly, cell survival remained above 90% even after 8 h of treatment, indicating negligible reduction in metabolic activity under these conditions. Based on these results, 100 µM HCQ was considered minimally cytotoxic and was therefore used in subsequent short-term experiments, including cell migration assays. To model wound repair, a ∼200 µm–wide scratch was introduced in a confluent monolayer of HeLa cells using a sterile pipette tip (**Figure 1A-i**).

**Figure 1:**
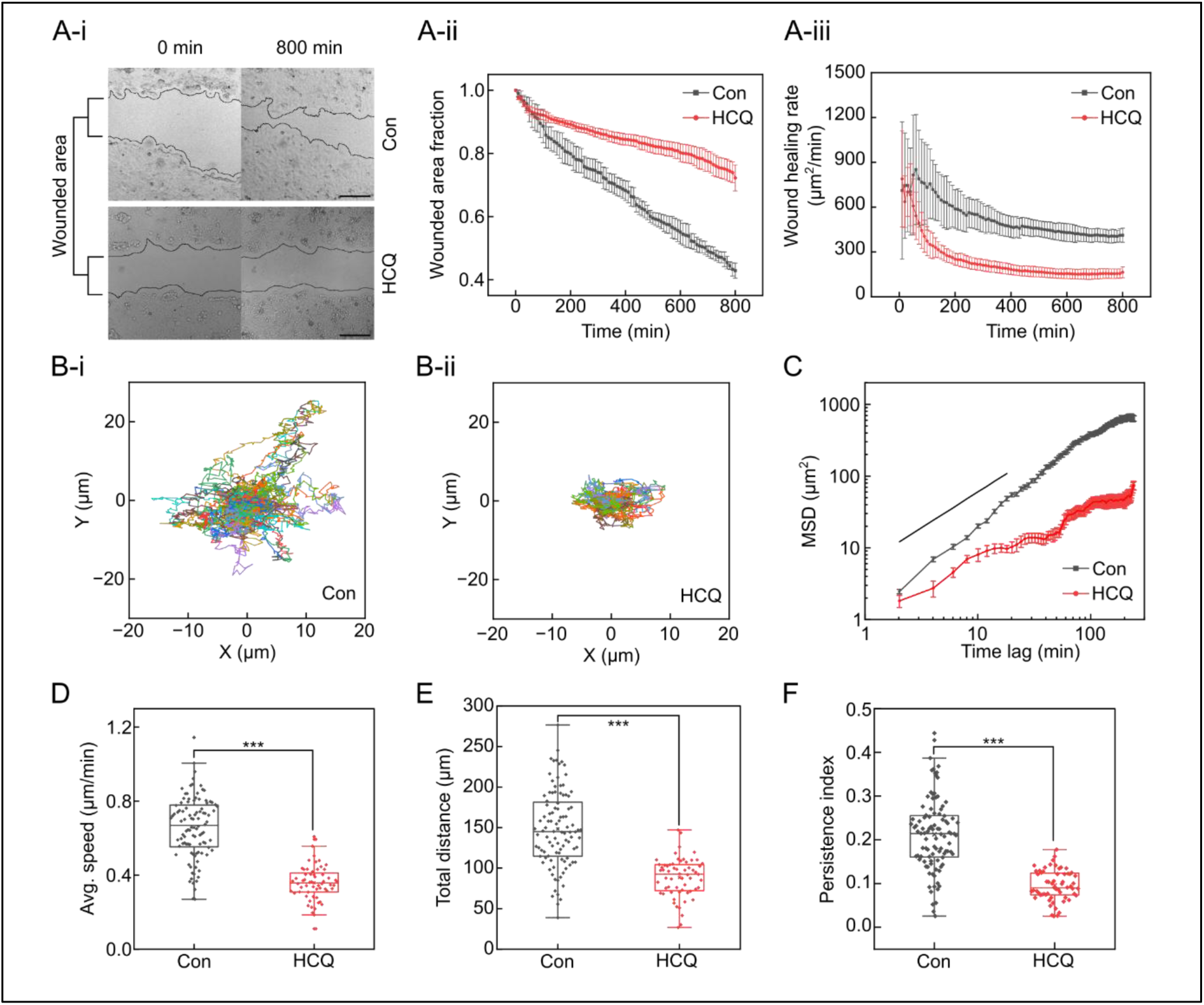
Hydroxychloroquine (HCQ) treatment impairs collective and single-cell migration. (A) Wound healing assay comparing control and HCQ-treated cells demonstrates markedly reduced migration with HCQ exposure. (A-i) Representative phase-contrast images at 0 and 800 minutes show noticeably slower wound closure in HCQ-treated cells (scale bar, 200 µm). (A-ii) Quantitative analysis of wound area fraction over time, and (A-iii) calculation of wound healing rates, both confirm impaired collective migration following HCQ treatment. (B) Single-cell migration assays display normalized trajectories under control conditions (B-i) and HCQ treatment (B-ii) tracked for 240 minutes, indicating reduced and less directed movement in the presence of HCQ. (C) Mean squared displacement (MSD) analysis reveals significantly lower displacement over time in HCQ-treated cells. (D–F) Boxplots summarizing average speed (D), total distance travelled (E), and randomness persistence index (displacement/distance) (F) illustrate that HCQ significantly decreases cellular speed and total motility, while increasing migratory randomness. Boxplots show median ± interquartile range; statistical comparisons performed using unpaired T-test (N_cells_>60; ***p < 0.001).

As expected, the wound area progressively decreased in control cells (**Figure 1A-ii, Video 1**). The rate and pattern of wound closure are influenced by parameters such as cell type, substrate coating, and assay conditions ^26^. While many scratch assays display an approximately linear closure over time, others exhibit nonlinear or biphasic kinetics ^26^. In our experiments, the wound closure rate in control HeLa cells was distinctly non–uniform (**Figure 1A-iii**), rising sharply from ∼700 µm²/min to ∼850 µm²/min within the first 100 minutes, followed by a gradual decline to ∼400 µm²/min during the subsequent 700 minutes. Intermittent spikes in recovery rate were observed in individual recovery traces (**Figure S1-B**), suggesting periodic bursts of collective migration. These findings indicate that HeLa cells display a time–dependent modulation of cell migration during wound closure.

We next assessed the effect of HCQ on wound healing. Treatment with 100 µM HCQ altered wound closure dynamics (**Figure 1A**). In contrast to control cells, which exhibited an early acceleration phase, HCQ-treated monolayers pre-incubated for 60 minutes prior to scratch formation failed to show the rapid increase in closure rate during the first 100 minutes. Instead, the closure rate decreased monotonically from ∼700 µm²/min immediately after scratch creation to ∼100 µm²/min over the following 800 min, indicating a progressive suppression of migratory activity. Moreover, the marked reduction in intermittent spikes in migration rate observed in HCQ-treated cells (**Figure S1-B**) suggest that HCQ compromises the ability of cells to generate coordinated, collective bursts of migration during wound repair.

A similar effect was observed in the wound healing assay performed with MDA-MB-231 cells, where representative images indicate that hydroxychloroquine (HCQ) treatment impairs wound closure over a period of 8 hours (**Figure S1-C**).

To discern whether HCQ impairs collective migration by affecting intercellular coordination or intrinsic single–cell motility, we examined isolated HeLa cells (**Video 2**). **Figure 1B** depicts 240 minutes trajectories of ≥25 control and HCQ-treated cells. Untreated cells displayed random migration with substantial displacement from their points of origin (**Figure 1B-i**). In contrast, cells pre-exposed to 100 µM HCQ exhibited markedly reduced motility, with most cells remaining confined close to their initial positions, indicating that HCQ treatment significantly suppresses cell movement (**Figure 1B-ii**). Quantitative analysis based on mean square displacement (MSD) confirmed a significant reduction in cell motility following HCQ treatment (**Figure 1C**). Both the average speed (**Figure 1D**) and total distance traversed (**Figure 1E**) were significantly diminished. The relationship between MSD and time (MSD = D·t^α^) yielded α ≈ 1 for control cells, indicating Brownian-like motion, whereas α < 1 for HCQ-treated cells suggested sub-diffusive confinement. Furthermore, the persistence index, PI (net displacement/total path length), was substantially lower following HCQ exposure (**Figure 1F**), indicating reduced directional persistence during migration.

Cell migration during wound healing depends critically on the actin cytoskeleton, which regulates lamellipodial protrusions through actin polymerization. Such protrusive activity is predominantly governed by Arp2/3–mediated nucleation of branched actin filaments ^27^. The observed retardation of wound healing by 100 µM HCQ (**Figure 1**) therefore implies that HCQ may perturb lamellipodia formation by interfering with actin polymerization dynamics.

Together, these results demonstrate that HCQ might substantially restrict both collective and individual cell migration by suppressing actin–driven protrusive activity. This prompted us to investigate whether HCQ induces reorganization of the cytoskeletal architecture, focusing specifically on actin and vimentin networks in the following analyses.

### HCQ Alters Actin and Vimentin Cytoskeletal Architecture

To elucidate the mechanism by which HCQ reduces cell migration, we next examined its effect on cytoskeletal organization in HeLa cells. Cells were divided into three spatial zones: edge (0 to 250 µm), inner (250 to 500 µm), and inner2 (>500 µm) to quantify spatial variations in cytoskeletal structure (**Figure 2A**). Comparative imaging of actin and vimentin filaments in control and HCQ–treated cells revealed that actin organization appeared largely similar upon visual inspection (**Figure 2B**). However, to detect subtle structural differences not apparent by eye, we computed the fractal dimension (FD), a quantitative metric of filament organization ^28^. Treatment with 100 µM HCQ for 1 hour selectively reduced actin filament density in cells located in the edge (**Figure 2C-i**).

**Figure 2:**
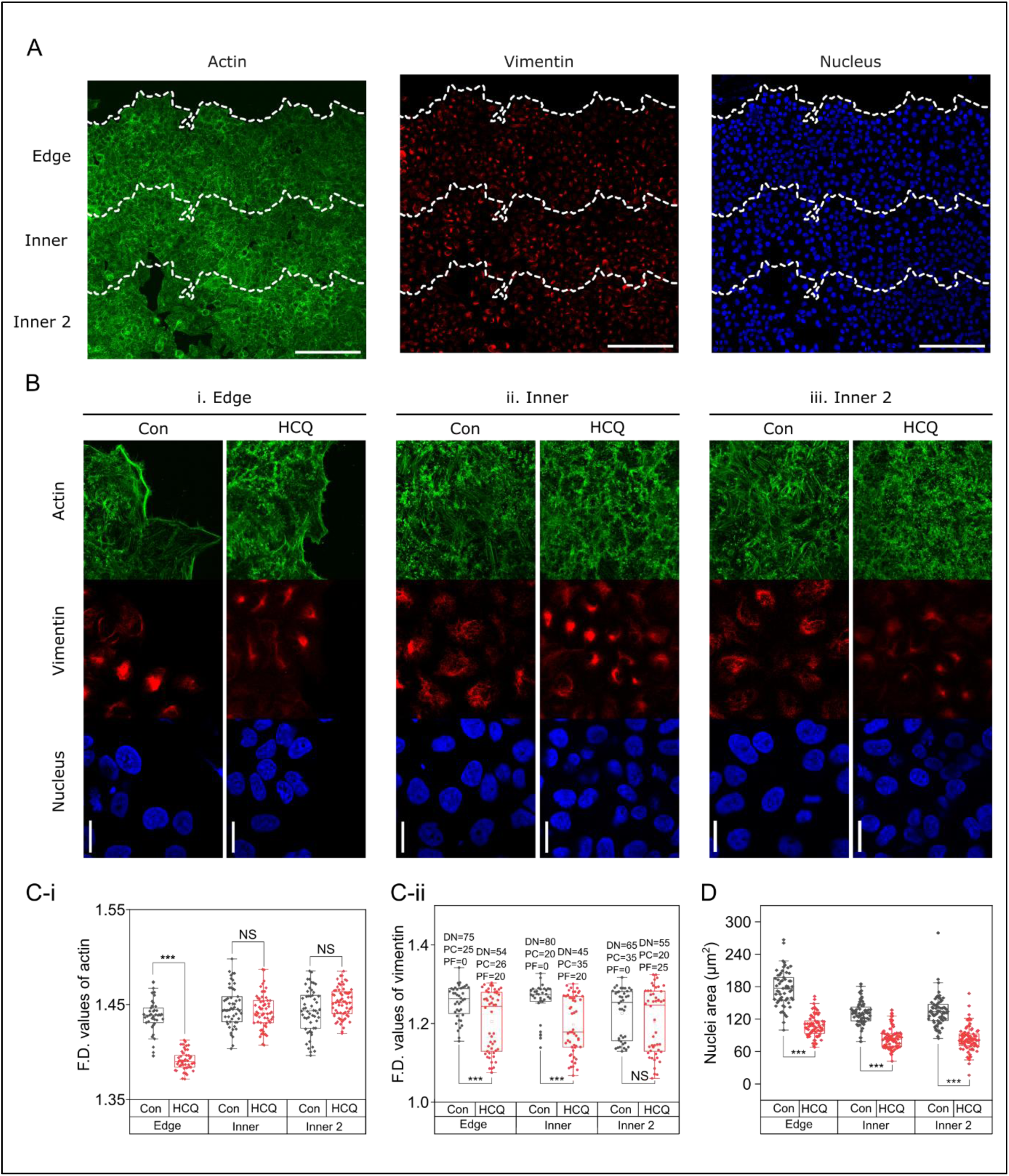
HCQ treatment results in a significant decrease in actin filament density at the wound edge while having minimal impact on the inner regions. (A) Representative low-magnification (10×) confocal images of control cells stained for actin, vimentin, and nuclei across the wound edge, inner, and inner 2 regions of the wound healing assay. (Scale bar, 200 µm) (B) High-magnification (63×) confocal images show actin (green), vimentin (red) and nuclei (blue)at the edge (i), inner (ii) and inner 2 (iii) regions with and without HCQ treatment.(Scale bar, 20 µm) (C) Quantification of actin (i) and vimentin (ii) filament density using fractal dimension (FD) analysis. Vimentin filament morphologies are categorized as dispersed network (DN), perinuclear cage (PC), and perinuclear foci (PF), and the percentages of these morphologies are presented in C-ii. (N_cells_ > 40, ***p < 0.001; NS, not significant, two-sample t-test) (D) Comparison of nuclear volume between control (Con) and hydroxychloroquine (HCQ) treated cells. (Ncells > 40, ***p < 0.001; NS, not significant, two-sample t-test)

Intermediate filaments, such as vimentin, provide mechanical stability, with vimentin deficient cells exhibiting a softer cytoplasm than wild type cells ^29^, and directional cues during collective migration and wound closure ^30^. In the majority of control cells (75%), vimentin filaments formed a dispersed cytoplasmic network (DN), characterized by filaments distributed throughout the cytoplasm without obvious perinuclear enrichment (**Figure S2**). In the remaining 25% of control cells, vimentin adopted a perinuclear cage (PC) organization, in which filaments formed a continuous or semi-continuous mesh surrounding the nucleus. Perinuclear foci (PF), defined as compact vimentin accumulations localized adjacent to the nuclear envelope, were not observed in control cells. Representative DN, PC, and PF morphologies and their corresponding FD values are shown in **Figure S2-A,B**. Upon HCQ treatment, vimentin organization was markedly altered. Under these conditions, 54% of cells retained a DN pattern, whereas 26% exhibited a PC partially enveloping the nucleus, and 20% displayed prominent PF formation (**Figure 2B, 2C-ii**). In addition, HCQ exposure led to a reduction in nuclear size in wounded cells (**Figure 2D**), which may be linked to changes in mechanical coupling between the vimentin intermediate filament network and the nucleus. We speculate that redistribution of vimentin toward perinuclear cages and foci could potentially alter mechanical connections between the cytoskeleton and the nucleus, thereby influencing nuclear morphology.

Together, these findings indicate that HCQ reorganizes cytoskeletal architecture by reducing actin filament density and altering vimentin distribution, particularly at the wound edge. Such remodelling may compromise the structural support and mechanical connectivity necessary for directed cell migration. Given these cytoskeletal changes, we next investigated whether HCQ also affects the dynamic lamellipodial protrusions-retraction activity that drive collective migration during wound closure.

### HCQ reduces lamellipodial dynamics and perturbs cytoskeletal remodelling

Because lamellipodial protrusion and retraction are key drivers of collective migration and coordination at the wound edge, the observed reduction in actin filaments prompted us to examine whether HCQ directly affects lamellipodial dynamics. Understanding this relationship is essential to link HCQ–induced cytoskeletal alterations to the impairment of collective migration during wound closure. To this end, we developed a quantitative framework to analyse lamellipodial protrusion–retraction activity, termed **Normalized Intensity Fluctuation Analysis** (**NIFA**) (**Figure 3A**). In this method, we recorded a 5-minute time lapse sequence, with an interval of 5 seconds between frames, and computed, for each pixel, the standard deviation (SD) of its temporal intensity fluctuations. Regions exhibiting active protrusive behaviour, such as lamellipodia, displayed higher SD values, generating a spatial map of protrusive activity. This approach enabled unbiased, high–resolution quantification of lamellipodial dynamics, independent of manual interpretation. In control wound cells, we observed multiple cycles of protrusion and retraction, whereas HCQ treatment markedly suppressed these lamellipodial fluctuations at the leading edge of the wound (**Figure 3A**). A comparable reduction in lamellipodial activity was also detected in isolated HCQ–treated cells (**Figure S3**).

**Figure 3.**
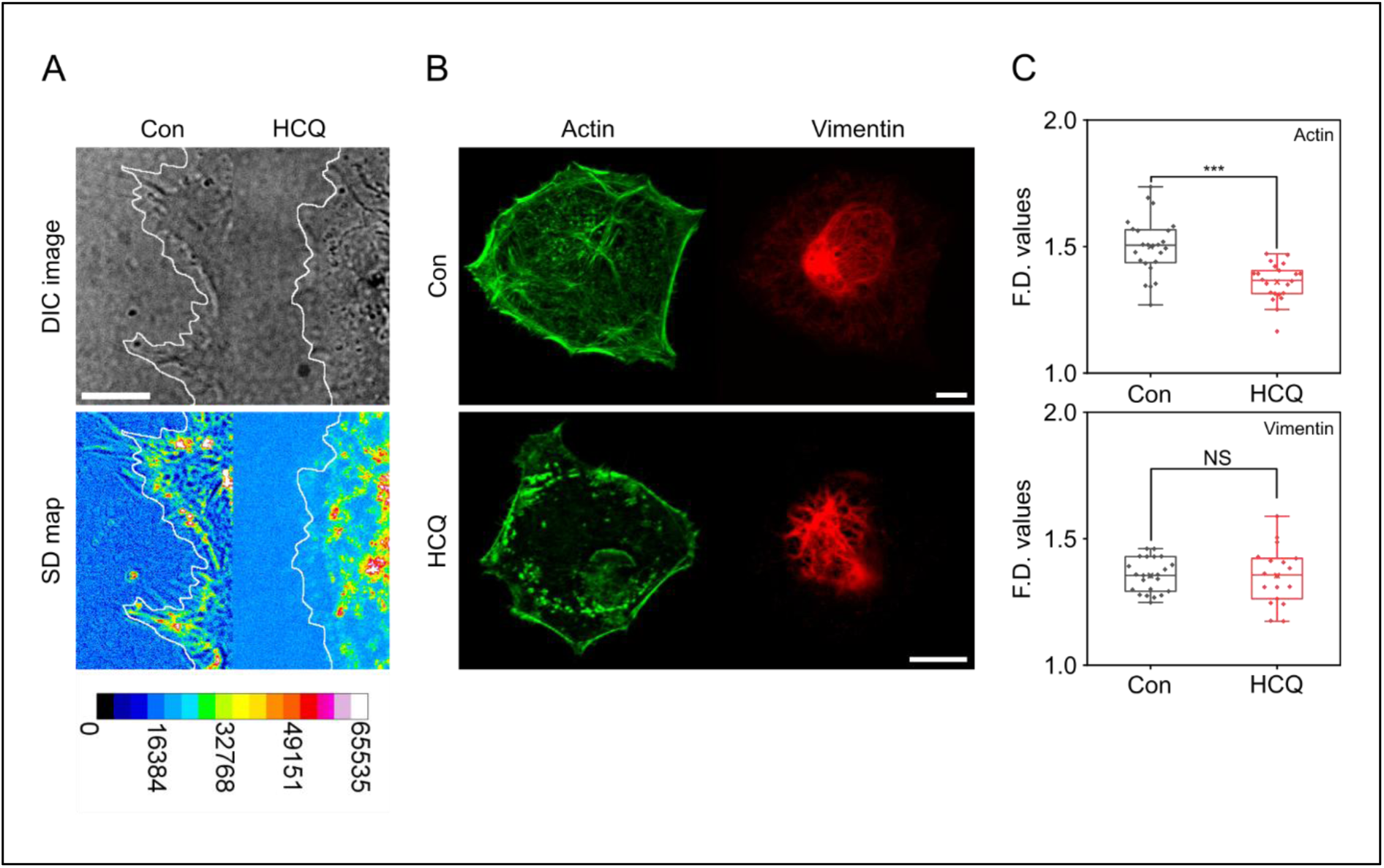
HCQ reduces lamellipodial dynamics and actin filament complexity in single cells. (**A**) Heatmaps showing lamellipodial protrusion–retraction activity in control and hydroxychloroquine (HCQ) treated cells reveal reduced edge dynamics upon HCQ treatment. (Scale bar, 10 µm) (**B**) Representative confocal images of single cells stained for actin and vimentin show disrupted actin organization in HCQ-treated cells, with minimal change in vimentin structure. Scale bar: 10 µm. (**C**) Quantification of filament complexity using fractal dimension (F.D.) analysis shows a significant decrease in actin F.D. in HCQ-treated cells, while vimentin F.D. remains unchanged. (N_cells_>20, ***p < 0.001; NS, not significant, two-sample *t*-test)

To determine whether the cytoskeletal effects of HCQ are specific to cues associated with collective migration or reflect a broader impact on filament organization, we analysed actin and vimentin architecture in isolated cells. In this setting, HCQ treatment disrupted the actin filament network, resulting in significant changes in FD values (**Figure 3B** and **3C**). Conversely, vimentin filaments were largely unaffected under the same conditions (**Figure 3B** and **3C**). Together with **Figure 2**—which show pronounced HCQ–induced reorganization of vimentin during wound healing—these results suggest that HCQ directly perturbs actin structure, whereas its effects on vimentin are context–dependent, differing between collectively migrating and isolated cells.

### HCQ induces cytoplasmic softening through cytoskeletal reorganization

Cytoplasmic softening is recognized as an early biophysical hallmark of apoptosis and can increase a cell’s susceptibility to cytotoxic agents ^31^. Given that HCQ treatment reorganizes both actin and vimentin filaments (**Figure 2, 3**), we hypothesized that these cytoskeletal changes may lead to cytoplasmic softening. To investigate this, we employed particle tracking micro-rheology (PTM) to measure intracellular stiffness. Unlike atomic force microscopy (AFM), which primarily probes cortical mechanics, PTM allows assessment of viscoelastic properties deep within the cytoplasm.

**Figure 4A** shows healthy cells embedded with fluorescently tagged microspheres ballistically delivered using a gene gun. **Figure 4B** presents two representative two-dimensional (2D) microsphere trajectories, tracked over 20 seconds, one from a control cell and one from a cell treated with HCQ. These trajectories were used to calculate the MSD for each particle (**Figure 4C**), which reflects the local viscoelastic properties of the surrounding cytoplasm. Because the cytoplasm is inherently heterogeneous, particles embedded in softer regions exhibit higher MSD values than those in stiffer regions. The observed variability among MSD curves confirms this mechanical heterogeneity. From MSD data, we computed the frequency–dependent complex modulus (G*), enabling extraction of both the elastic modulus, G′ (**Figure 4D**) and the viscous modulus, G’’ ^32^. The dynamic viscosity, η, was calculated by dividing the viscous modulus, G’’, by the corresponding angular frequency, ω (**Figure 4E**).

**Figure 4:**
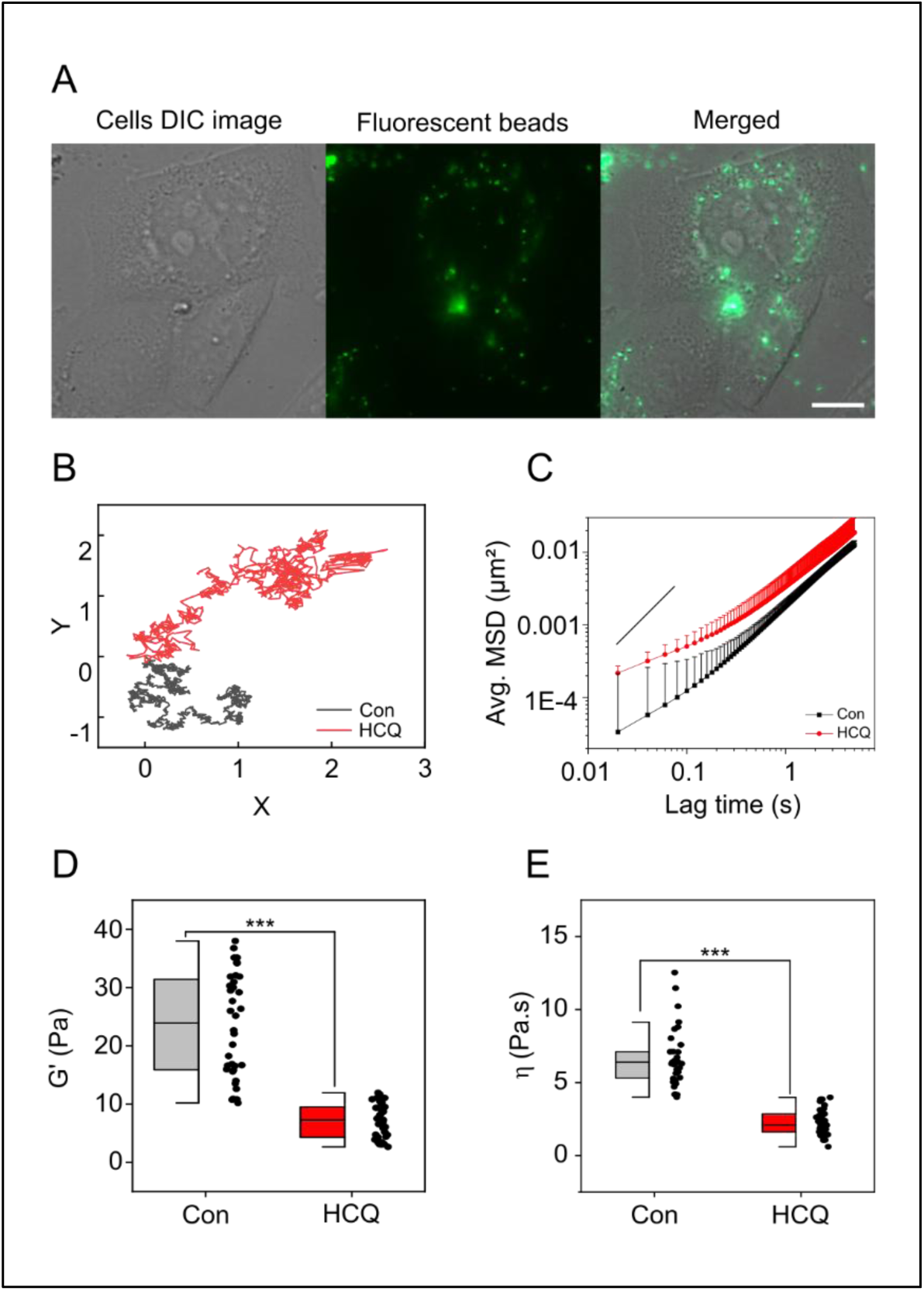
HCQ-treatment results in overall softening of the cytoplasm. (A) Localization of fluorescently labelled beads (green) inside HeLa cells for particle tracking micro-rheology. Scale bar, 10 µm (B) Representative trajectories of beads embedded in the cytoplasm of control and hydroxychloroquine (HCQ)-treated cells. (C) Average mean squared displacement (MSD) comparison between control and HCQ-treated cells. Comparison of elastic modulus, G’ (D) and dynamic viscosity, η (E) between control and HCQ-treated cells. (N_cells_>30, ***p < 0.001; Mann-Whitney U test)

Despite the inherent variability in particle motion, a consistent increase in MSD values following HCQ exposure indicated substantial cytoplasmic softening. Correspondingly, the average elastic modulus decreased, and mechanical heterogeneity was reduced. Moreover, HCQ–treated cells displayed a decline in cytoplasmic viscosity, together with a more uniform viscous behaviour across the population. These results collectively demonstrate that HCQ treatment renders the cytoplasm softer and mechanically homogeneous. This softening likely arises from cytoskeletal reorganization and may sensitize cells to subsequent cytotoxic stress.

### HCQ Binds Cooperatively to Actin and Perturbs Its Secondary Structure

To determine whether these biomechanical effects originate from a direct molecular interaction between HCQ and actin, we next analysed their binding characteristics in vitro. **Figure 5A** displays the fluorescence emission spectra of actin (10 µM) excited at 260 nm, recorded in the absence and presence of increasing concentrations of HCQ. The intrinsic fluorescence of actin, observed between 300 and 350 nm, progressively decreased upon HCQ addition up to ∼9 µM, after which the intensity plateaued. Concurrently, an additional emission feature between 360 and 450 nm, characteristic of HCQ, increased proportionally with its concentration, indicating that HCQ binding results in fluorescence quenching of actin. Time–resolved fluorescence measurements revealed a marked reduction in the mean lifetime (**Figure 5B** and **5C, Figure S4**) of actin upon HCQ addition, signifying alterations in the microenvironment of aromatic residues (tryptophan, tyrosine, and phenylalanine). To quantify the binding affinity, the fluorescence change of actin with respect to HCQ concentration was fitted using the Hill equation, yielding a dissociation constant (Kd) of ∼3.6 ± 0.1 µM (**Figure 5D**). Complementary experiments, in which HCQ (10 µM) was directly excited in the presence of varying actin concentrations, produced a comparable Kd (∼4.0 ± 0.4 µM; **Figure S4**), consistent with the initial measurement. These values fall within the range reported for small–molecule ligands interacting with actin ^33^. Notably, the Hill coefficient (n) exceeded 1 in both analyses, suggesting cooperative binding—wherein HCQ engages multiple sites on actin and the binding of one molecule enhances subsequent interactions.

**Figure 5:**
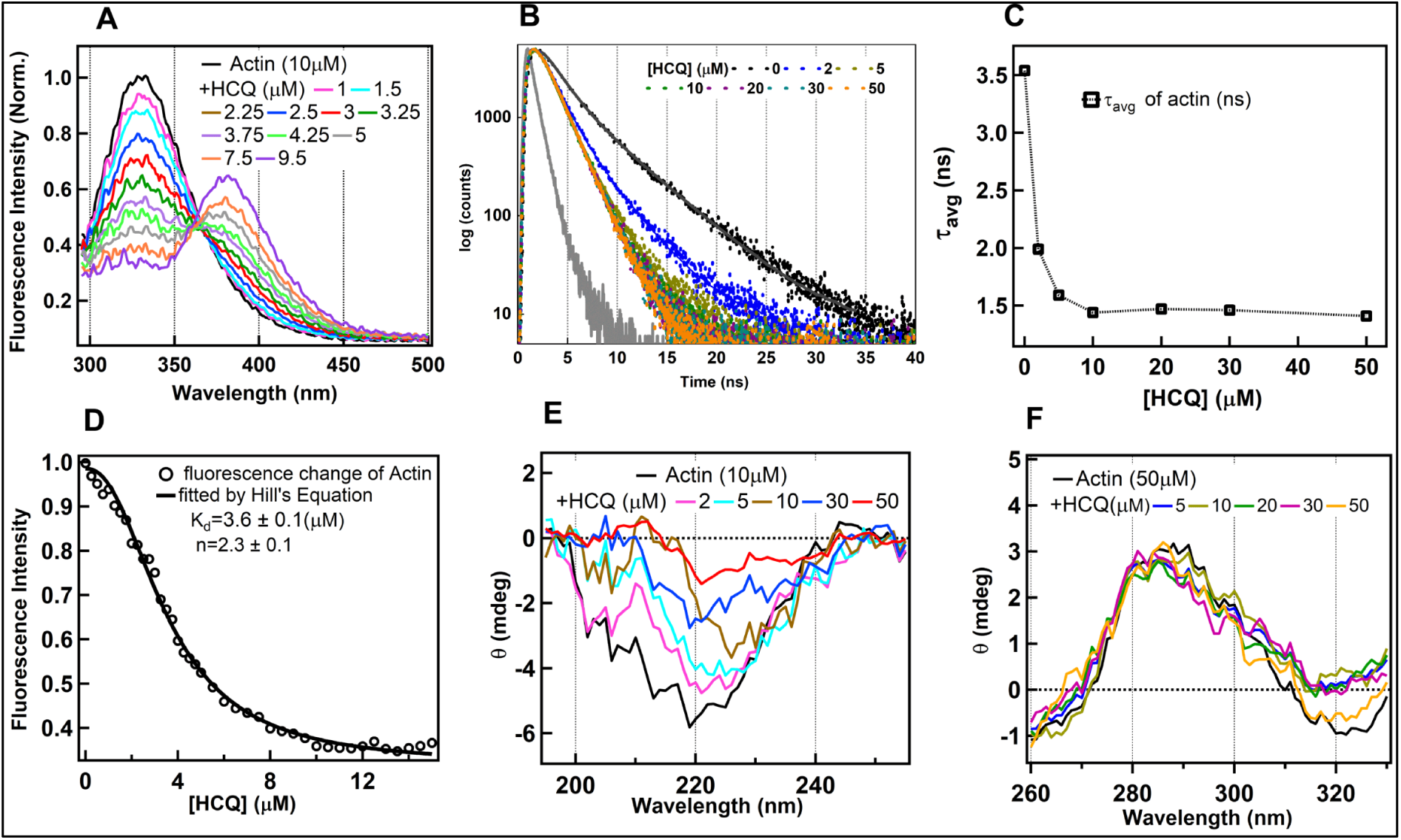
(A) The fluorescence emission spectra of actin (10 µM) in in the absence and presence of the different concentrations of HCQ obtained on the excitation at 280 nm. The spectral feature in the 300-350 nm with the peak maxima (330 nm) belongs to the actin whereas the peak feature in the 350-450 nm with the peak maxima (380 nm) corresponds to the HCQ. (B) The lifetime traces of actin (10 µM) in the absence and presence of the different concentrations of HCQ obtained on the excitation at 280 nm. (C) The trend of the change in the average life time (ns) of actin with the addition of different concentrations of HCQ. (D) The plot of the change in the emission maxima of actin (∼330 nm) with the addition of the HCQ. The K_d_ value of the binding of actin with HCQ was obtained using the Hill’s equation. (E)The change in the CD spectra representing the secondary structure of actin (10 µM) with the addition of HCQ. (F) The change in the CD spectra representing the tertiary structure of actin (50 µM) with the addition of HCQ.

To determine how HCQ binding affects actin conformation, circular dichroism (CD) spectra were acquired in both far–UV and near–UV regions, corresponding respectively to secondary and tertiary protein structure (**Figure 5E and 5F**). In the far–UV region, a progressive reduction in ellipticity was observed as HCQ concentration increased, indicating perturbation of actin’s α–helical and β–sheet content. Such changes are reminiscent of structural disruptions caused by classical protein denaturants ^34^. By contrast, the near–UV CD spectra remained largely unaltered, implying minimal influence on tertiary aromatic and disulfide–bond configurations. Taken together, these findings indicate that HCQ induces partial unfolding of G–actin predominantly through perturbations in its secondary structure.

Overall, these observations establish that HCQ binds cooperatively to actin with micromolar affinity, altering its secondary structural integrity without significantly disturbing tertiary organization. This cooperative binding may underlie the compound’s ability to disrupt actin polymerization and ABPs (actin binding protein) ability to bind actin filaments, as explored in subsequent analyses.

### HCQ reduces actin polymerization rate and impairs interactions between actin and actin binding proteins

To further assess how HCQ influences actin polymerization dynamics, we examined its effect on actin turnover in vitro using pyrene–labelled actin. Pyrene fluorescence increases markedly as monomeric actin polymerizes into filamentous (F–actin) structures ^35^. **Figure 6A** compares the polymerization kinetics of actin under control and HCQ–treated conditions. Exposure to 100 µM HCQ altered polymerization behaviour. The fluorescence traces were fitted to a kinetic model (A·(1−e^−kx^)), yielding low and uniform residuals (**Figure 6A**), confirming that the data were well described by the model. **Figure 6B** presents polymerization rates (k) derived from three independent measurements, demonstrating that 100 µM HCQ reduces the actin polymerization rate *in vitro*.

**Figure 6:**
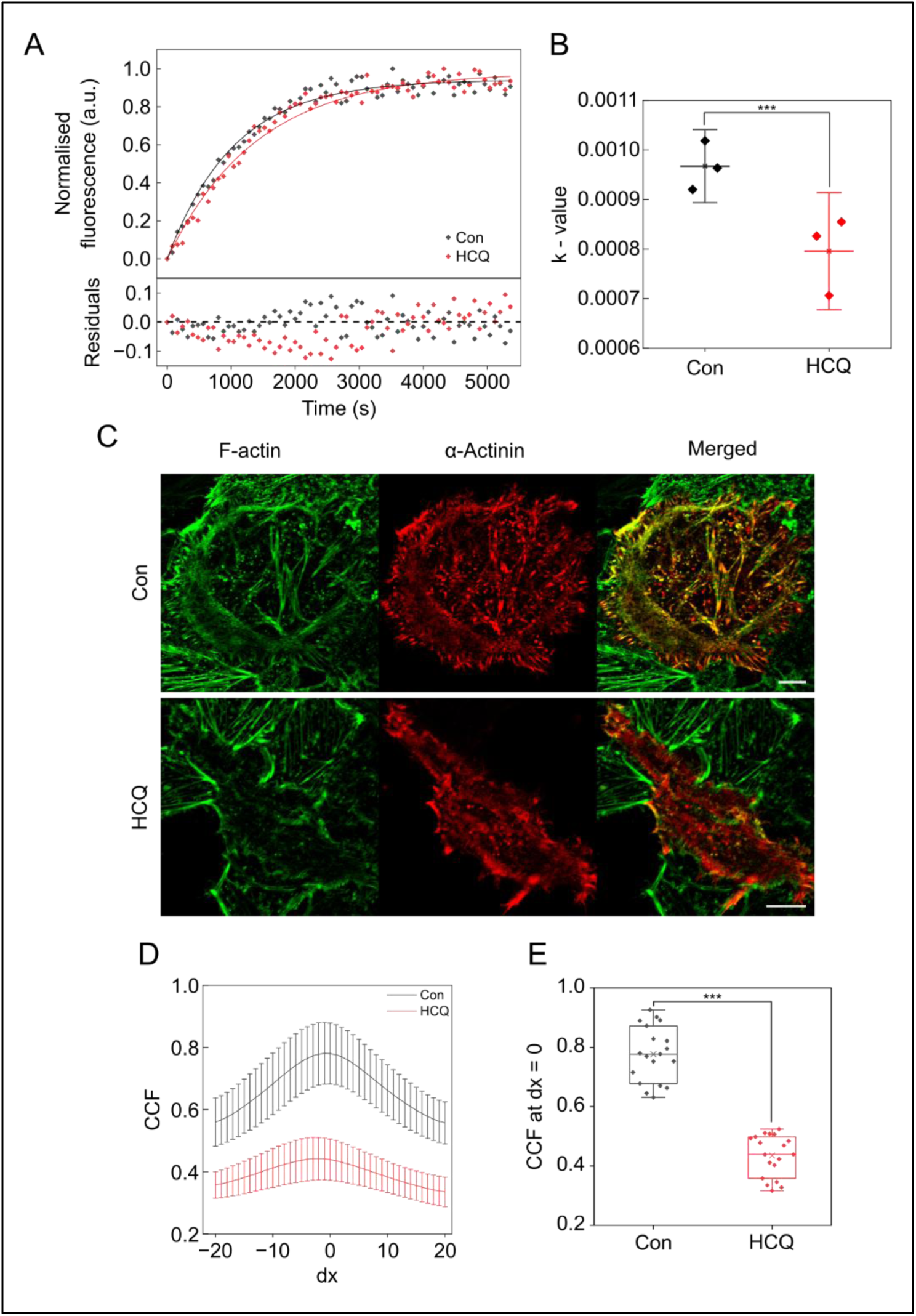
Hydroxychloroquine (HCQ) reduces colocalization between α-actinin and F-actin. (A) Pyrene–actin polymerization kinetics in the absence and presence of HCQ. Scatter points represent normalised fluorescence intensity data, and solid lines denote fitted polymerization curves. The bottom section of the graph represents the residuals for the fitting. (B) Rate constant (*k*) derived from the fitted polymerization curves, presented as mean ± SD for control (Con) and HCQ-treated samples. (C) Representative confocal images of cells expressing mCherry-α-actinin (red) and stained with AF488–phalloidin for F-actin (green) under control (Con) conditions (top) and after treatment with HCQ (bottom). (Scale bar, 20 µm) (D) Cross-correlation function (CCF) plots showing the spatial relationship between α-actinin and F-actin with an x-shift of ±20 pixels. (E) Box plot of CCF values at dx = 0 demonstrating a significant reduction in colocalization between α-actinin and F-actin upon HCQ treatment (N_cells_ > 15, ***p < 0.001, unpaired t-test)

Given the observed alterations in actin structure, we next examined whether HCQ affects the interaction of ABPs (actin binding proteins) with filamentous actin. As a model ABP, we analysed α–actinin, a crosslinking protein that binds F–actin to stabilize cytoskeletal architecture. HeLa cells were transiently transfected with mCherry-α-actinin, and F–actin was visualized using phalloidin tagged with Alexa Fluor 488 (AF488) (**Figure 6C**). In control cells, mCherry-α-actinin exhibited strong colocalization with F–actin. However, treatment with 100 µM HCQ visibly decreased this overlap in the representative images (**Figure 6C**). To quantify colocalization, we computed Van Steensel’s cross–correlation function (CCF) along the x–axis between mCherry-α-actinin and phalloidin–AF488 fluorescence (**Figure 6D**). In untreated cells, CCF values peaked at zero displacement, consistent with strong spatial colocalization, whereas HCQ treatment reduced the peak CCF (dx = 0) value (**Figure 6E**). This reduction reflects a loss of colocalization between α–actinin and F–actin filaments.

Together, these findings demonstrate that HCQ slows actin polymerization kinetics and disrupts the binding of actin–associated proteins such as α–actinin. By perturbing both actin dynamics and ABP interactions, HCQ may broadly impair cytoskeletal integrity and cellular motility.

## Discussion

Actin and microtubule networks are central to cancer progression because they control key processes such as the cell cycle, cell shape regulation, and migration. While microtubule-targeting agents like vinca alkaloids and taxanes are already well integrated into clinical treatment ^36^, therapeutics that directly target the actin cytoskeleton remain largely undeveloped and distant from clinical use. One major reason for the lack of actin-targeting anticancer agents is the limited molecular-level understanding of how drugs interact with the actin cytoskeleton and how these interactions influence downstream biological responses.

HCQ, widely known for its immunomodulatory and antiviral effects, has recently been proposed as a potential anticancer agent in several studies ^25,37^. In this study, we focus on elucidating how HCQ affects the cytoskeleton of cancer cell lines and uncover the molecular mechanisms underlying these effects. The findings indicate that HCQ impairs collective cell migration by affecting actin dynamics and disrupting cytoskeletal integrity at multiple structural and functional levels. Together, our results present a mechanistic framework linking the outcome of HCQ-actin interaction in cell motility, cytoskeletal remodelling, and intracellular mechanics.

HCQ significantly attenuated wound closure during collective migration, abolishing the periodic bursts of motile activity typically observed in coordinated cell movement. Single-cell tracking revealed that this suppression was accompanied by a reduction in overall motility, directional persistence, and displacement, establishing that HCQ affects not only collective coordination but also the intrinsic migratory capacity of individual cells. The loss of synchronized migration suggests that HCQ interferes with cytoskeletal processes critical for the generation and propagation of mechanical forces across cell–cell junctions.

Cytoskeletal imaging and quantitative analysis further revealed that HCQ alters both actin and vimentin organization, particularly in cells located at the wound edge, where actin polymerization and vimentin alignment are essential for front–rear polarity. Treatment with HCQ reduced actin filament density and induced a striking redistribution of vimentin from a dispersed cytoplasmic network to perinuclear aggregates, indicating broad cytoskeletal remodelling. Consistent with these structural alterations, lamellipodial dynamics, assessed through Normalized Intensity Fluctuation Analysis, were markedly diminished following HCQ treatment, linking reduced migratory behaviour to impaired actin-driven protrusive activity.

The impact of HCQ on cytoskeletal organization extended to cellular mechanics. Particle tracking micro-rheology revealed a substantial increase in mean square displacement of intracellular probes, accompanied by a reduction in both elastic modulus and viscosity. These measurements indicate that HCQ induces cytoplasmic softening and mechanical homogenization, likely resulting from actin depolymerization and cytoskeletal disassembly. Cytoplasmic softening is a well-recognized biophysical hallmark of early apoptosis and can enhance susceptibility to cytotoxic stimuli, suggesting that HCQ-induced mechanical changes may contribute to its reported anti-proliferative and sensitizing effects in cancer cells.

At the molecular level, steady-state and time-resolved fluorescence analyses established that HCQ binds directly to actin with micromolar affinity. The binding was cooperative, as reflected by a Hill coefficient greater than one, implying multiple interacting sites that reinforce each other’s affinity. Circular dichroism spectroscopy revealed that HCQ binding disrupts actin’s α-helical and β-sheet content while sparing tertiary aromatic and disulfide configurations, consistent with partial unfolding of the monomer. These conformational perturbations likely hinder filament assembly. Indeed, pyrene-actin assays confirmed that HCQ reduces the rate of actin polymerization, while cellular colocalization analysis showed that the drug weakens the interaction between filamentous actin and α-actinin, an essential actin-binding protein that stabilizes stress fibres and lamellipodia.

Together, these findings present a coherent mechanistic view: HCQ directly associates with actin, inducing subtle structural distortions that impair actin polymerization and weaken interactions with actin-binding partners. The resulting cytoskeletal destabilization manifests as reduced lamellipodial activity, altered vimentin organization, cytoplasmic softening, and loss of coordinated cell migration. By modulating actin architecture and the mechanical properties of cells, HCQ effectively suppresses the cytoskeletal machinery essential for cell motility. This study thus provides mechanistic insight into how HCQ functions as a biophysical modulator of cytoskeletal integrity, offering a molecular explanation for its reported anti-migratory and anti-metastatic properties in cancer and potentially other pathophysiological contexts.

## Material and methods

### Cell culture

HeLa and MDA-MB-231 cells were obtained from the National Centre for Cell Science (NCCS), Pune, India. Cells were maintained in Dulbecco’s Modified Eagle Medium (DMEM; HiMedia, #AL007G) supplemented with 10% foetal bovine serum (FBS; HiMedia, #RM10434) and 5% penicillin–streptomycin antibiotic cocktail (HiMedia, #A001A) to prevent bacterial contamination. Cultures were incubated at 37 °C in a humidified atmosphere containing 5% CO₂. Cells were passaged upon reaching ∼70% confluence to ensure optimal growth and proliferation, and were routinely tested for mycoplasma contamination. For imaging experiments, cells were seeded on Poly-L-lysine–coated confocal dishes to promote optimal adhesion. During live-cell imaging experiments, cells were maintained in Leibovitz’s L-15 medium (HiMedia) instead of DMEM to eliminate the requirement for CO₂ incubation.

### MTT assay for cell viability

Cell viability following hydroxychloroquine (HCQ) treatment was assessed using the MTT assay. HeLa cells were seeded in a 96-well plate and allowed to adhere overnight. Cells were then treated with 100 µM HCQ for 8, 16, and 24 hours. Following treatment, MTT solution was added to each well and incubated for 3 hours at 37 °C to allow the formation of formazan crystals. The medium was subsequently removed and the crystals were dissolved in dimethyl sulfoxide (DMSO). Absorbance was measured at 570 nm using a microplate reader (Molecular Devices SpectraMax iD5). Cell survival was calculated relative to untreated control cells at 0 hour, which were normalized to 100%.

### Drug treatment

Hydroxychloroquine sulphate (HCQ; Tocris®, #5648) was dissolved in nuclease-free, molecular biology–grade double-distilled water to prepare a 5 mM stock solution, which was stored at –20 °C. For treatments, HCQ was added to the culture medium at a final concentration of 100 µM for 1 h, unless otherwise specified.

### Wound healing assay and analysis

For the wound healing assay, cells were seeded onto 35-mm glass-bottom confocal dishes two days prior to the experiment and allowed to proliferate until reaching 100% confluence. On the day of the assay, hydroxychloroquine (HCQ) was added to the experimental group at a final concentration of 100 µM, 1 h before scratching. A linear wound was then introduced at the centre of the confluent monolayer using a 10 µL pipette tip. Following scratching, the culture medium was removed, and the dishes were gently washed with 1× PBS to eliminate detached cells. Fresh Leibovitz’s L-15 medium (without foetal bovine serum) was then added, and the dishes were maintained on a heated microscope stage at 37 °C. For HCQ-treated groups, Leibovitz’s L-15 medium was supplemented with HCQ to a final concentration of 100 µM for the entire duration of the experiment. Time-lapse imaging of the wounded regions was performed for 800 minutes at 10-minutes intervals. The acquired images were analysed in **Fiji®** using the *Wound Healing Size Tool* plugin to quantify the wound area fraction over time and to calculate the wound healing rate. The wound healing rate (WHR) was determined using the following equation:

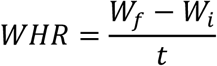

where W_i_ and W_f_ represent the initial and final wound areas, respectively, and t is the duration of the experiment.

### Single Cell Tracking

For single-cell tracking, nuclei were stained with Hoechst 33342 (Thermo, #62249). Cell movements were recorded as time-lapse sequences for 240 minutes with a 2-minutes interval between successive frames. Images were acquired in two channels—DIC and DAPI (excitation wavelength ∼330–380 nm)—to visualize both the nucleus and cell boundaries. Cell trajectories were analysed using the *TrackMate* plugin in ImageJ, which detected the x–y coordinates of nuclear centroids at each time point. From these coordinates, trajectories of individual cells were reconstructed. Mean squared displacement (MSD), average migration speed, total distance travelled, and persistence index were calculated from the trajectories using a custom MATLAB® script (available upon request).

### Cell fixation, permeabilization and immunostaining

Cells were fixed with 4% paraformaldehyde (HiMedia, #TC703) in 1× PBS for 20 minutes at room temperature, followed by three washes with 1× PBS. Permeabilization was performed using 0.1% Triton X-100 (Sigma-Aldrich, #X100) in 1× PBS for 3 minutes. Non-specific binding was blocked with 2% bovine serum albumin (BSA; HiMedia, #MB083) in 1× PBS for 1 h at room temperature. Actin filaments were stained with Alexa Fluor® 488–conjugated phalloidin (Invitrogen, #A12379; 1:200 in 1× PBS) for 30 minutes at room temperature in the dark. For vimentin immunostaining, cells were incubated with an anti-vimentin rabbit monoclonal primary antibody (Cell Signaling Technology, #5741; 1:200 in 1× PBS with 2% BSA) for 16 hours at 4 °C. The cells were then washed three times with 1× PBS and subsequently incubated with an Alexa Fluor 594–conjugated anti-rabbit secondary antibody (Cell Signaling Technology, #8889; 1:500 in 1× PBS with 2% BSA) for 1 hour at room temperature in the dark. Finally, three additional washes with 1× PBS were performed to remove unbound secondary antibody. Nuclei were counterstained using Hoechst 33342 (Thermo, #62249; 1 µg/mL in 1× PBS) for 10 minutes at room temperature in the dark and washed once with 1× PBS. Samples were maintained in 1× PBS, and stored at 4 °C until imaging. The same protocol was used to fix and stain actin and vimentin filaments in cells during the wound healing assay, with fixation performed two hours after scratch formation.

### Microscopy and image acquisition

Immunofluorescence images of actin, vimentin, and nuclei were acquired using a Zeiss LSM 880 confocal microscope equipped with a 10x and 63× oil immersion objective. Collective cell migration during the wound healing assay and single-cell migration experiments were imaged using a Zeiss AXIO Observer.Z1 wide-field epifluorescence microscope equipped with a Hamamatsu ORCA Flash 4.0 CMOS camera and a 10× objective. Image acquisition was performed using either ZEN 2.0 (Zeiss) or μManager software ^38^.

### Fractal dimension analysis

Fractal dimension (FD) values of actin and vimentin filaments were calculated by the differential box counting (DBC) method available in the *FracLac* plugin (v2.5) for ImageJ ^39^. 16-bit grayscale images were used for the analysis. To minimize background noise and ensure accurate FD calculation, background pixel values were converted to zero. Imaging parameters were kept constant across all samples to allow reliable comparisons.

### Particle tracking micro-rheology (PTM)

The entire procedure of PTM was performed following standard protocols ^40,41^. Fluorescently tagged carboxylate-modified microspheres (200 nm diameter; Invitrogen, #F8811) were introduced into cells by ballistic delivery using a gene gun system (Helios® Gene Gun, Bio-Rad). After delivery, cells were washed with 1× PBS to remove unbound beads and detached cells, and subsequently incubated under standard culture conditions (37 °C, 5% CO₂) in serum-free DMEM medium for 16 hours to recover from the microsphere delivery procedure. Cells containing fluorescent microspheres were then positioned on a microscope stage maintained at 37 °C. Microsphere movements were recorded as movies using a high-speed Hamamatsu Orca Flash 4.0 CMOS camera controlled with μManager software ^38^. Each recording consisted of 1000 frames, corresponding to a total duration of 20 seconds, with an exposure time of 20 milliseconds per frame. Trajectories of individual microspheres were extracted using the MOSAICsuite plugin for ImageJ ^42^. At least 5–10 beads per cell were tracked to quantify the rheological properties of the cytoplasm. Mean squared displacement (MSD) values were calculated from the x and y coordinates of these trajectories. Out of the 1000 frames, only the first 250 frames were considered for MSD calculations to eradicate potential errors at later time points. The frequency dependent elastic (G’) and the viscous (G’’) moduli were algebraically derived from these MSD values ^32^. Dynamic viscosity (η) was computed by dividing viscous moduli (G’’) by the corresponding angular frequency, ω.

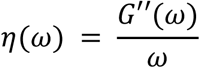

A custom MATLAB script was used for computing the MSD, elastic modulus (G’) and dynamic viscosity (𝜂) values.

### Pyrene actin polymerization assay

Pyrene-labelled muscle actin (Cytoskeleton, #AP05) was used to monitor actin polymerization kinetics. Actin was first diluted to 0.45 mg/mL in General Actin Buffer (5 mM Tris-HCl pH 8.0, 0.2 mM CaCl₂) supplemented with 0.2 mM ATP and 1 mM DTT, and incubated on ice for 1 hour to depolymerize residual oligomers. The solution was then centrifuged at 14,000 rpm for 30 minutes at 4 °C to remove residual nucleating centres. Polymerization assays were performed in two experimental conditions: (1) actin alone (untreated), (2) actin in the presence of 100 μM HCQ. For each condition, 200 μl of actin solution was pipetted into duplicate wells of a clear bottom 96-well plate. Wells containing only buffer without actin were used to establish background fluorescence. Baseline fluorescence was recorded for 3 minutes using a fluorescence spectrophotometer (excitation 360 nm, emission 405 nm; Molecular Devices SpectraMax iD5). Polymerization was initiated by adding 20 μL of 10× Actin Polymerization Buffer (500 mM KCl, 20 mM MgCl₂, 10 mM ATP) to each well, followed by gentle mixing. Fluorescence readings were acquired every 30 seconds for 2 hours. Fluorescence enhancement, which reflects actin filament formation, was analysed to generate polymerization curves for each experimental condition.

### Lamellipodial activity analysis

Lamellipodial dynamics were analysed in cells located at the leading edge of the wound. Time-lapse images were acquired using a Zeiss AXIO Observer.Z1 wide-field epifluorescence microscope with a 63× oil immersion objective. Image sequences were captured for 10 minutes with a 10-second interval between frames. Lamellipodial activity was quantified using a MATLAB® script by computing the standard deviation of pixel intensity values within the lamellipodium over time, which served as an indicator of protrusion and retraction dynamics.

### Biophysical Characterization of HCQ-actin interaction

The fluorescence of actin (10 µM) was recorded in the absence and presence of increasing concentrations of HCQ using a Fluoromax-4 spectrometer. Actin was excited at 280 nm, and emission spectra were collected until no change in fluorescence maximum was observed upon HCQ addition. Fluorescence lifetime measurements of actin (10 µM), both without and with varying concentrations of HCQ, were performed using a 280 nm picosecond diode laser as the excitation source. Emission was collected at 330 nm under magic-angle polarization and detected using a multichannel photomultiplier. The instrument response function (IRF), determined using a liquid scatterer, exhibited a typical full width at half maximum (FWHM) of ∼400 ps. The fluorescence decay curves were analyzed by fitting to a multiexponential function: I(t) = Σ Bi exp(−t/τ_i_), where 𝜏_𝑖_represents the fluorescence lifetime of each component and 𝐵_𝑖_is the corresponding pre-exponential factor. The decay profiles were deconvoluted with the instrument response function. The quality of the fit was assessed using the reduced chi-square (χ²) value and by examining the distribution of weighted residuals. The χ² values were close to 1, and the residuals were randomly distributed around zero, indicating an excellent fit. The binding affinity of HCQ to actin was determined by plotting the change in actin fluorescence intensity as a function of HCQ concentration and fitting the data using the Hill equation given below.

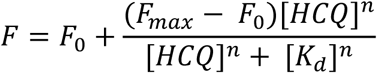

Here, F_0_ and F represent the fluorescence intensity of the actin in the absence and presence of HCQ whereas F_max_ refers to the case in which the fluorescence intensity of actin does not change further with the addition of HCQ. The value of 𝑛 depict the Hill coefficient reflecting binding stoichiometry/cooperativity. Additionally, the dissociation constant for the actin–HCQ interaction was independently estimated by monitoring the fluorescence of HCQ (10 µM), excited selectively at 330 nm, in the absence and presence of increasing concentrations of actin until saturation was achieved.

To investigate possible structural changes in actin upon HCQ binding, circular dichroism (CD) spectra were recorded using a JASCO J-1500 spectropolarimeter at 22 ± 2 °C. Measurements were carried out in the far-UV region (10 µM actin) to assess secondary structure and in the near-UV region (50 µM actin) to examine tertiary structural changes, both in the absence and presence of varying concentrations of HCQ.

### **T**ransfection of α-actinin

For transient expression of α-actinin, cells in each 35 mm dish were transfected with 2.5 µg α-actinin–mCherry plasmid (Addgene, #54975) using 15 µL DOTAP Liposomal Transfection Reagent (Sigma-Aldrich, #11202375001). Plasmid and DOTAP were mixed in serum-free DMEM according to the manufacturer’s instructions, incubated for 10 minutes at room temperature, and added to the cells. Following 6 hours of incubation, the transfection medium was replaced with complete DMEM. Cells were treated with drugs and fixed 48 hours post-transfection.

### Colocalization analysis of actin and α-actinin

Cells transfected with α-actinin–mCherry plasmid were fixed and subsequently stained with phalloidin–Alexa Fluor 488 to visualize actin filaments. Colocalization between α-actinin–mCherry and actin filaments was quantified using the JACoP plugin (Just Another Colocalization Plugin, version 2.1.4, 21/02/03) for ImageJ ^43^. Cross-correlation function (CCF) analysis with an x-shift of 20 pixels was performed to evaluate the spatial relationship between the two fluorophores.

## Funding

**Deepak Kumar Sinha** was supported by grants from the Department of Science and Technology, Ministry of Science and Technology, India (DST; Grant No. SB/S0/BB-101/2013 and CRG/2022/005356), the Department of Biotechnology, Ministry of Science and Technology, India (DBT; Grant No. BT/PR6995/BRB/10/1140/2012), and the Indian Association for the Cultivation of Science (IACS). **Dipanjan Ray** was supported by a fellowship from the Council of Scientific and Industrial Research (CSIR), India (File No. 09/080(1134)/2019-EMR-I), and by institutional funding from IACS. **Asim Bisoi** was supported by fellowship received from DST INSPIRE. **Bipasa Chaudhuri** was supported by a fellowship from the University Grants Commission (UGC), India.

## Author contributions

**Dipanjan Ray:** Investigation, Formal Analysis, Visualization, Software, Writing-Reviewing and Editing. **Asim Bisoi:** Investigation, Formal Analysis, Visualization, Writing-Reviewing and Editing. **Bipasa Chaudhuri:** Investigation, Formal Analysis, Visualization, Writing-Reviewing and Editing. **Prashant Chandra Singh:** Conceptualization, Supervision, Funding Acquisition, Writing-Original draft preparation, Writing-Reviewing and Editing. **Deepak Kumar Sinha:** Conceptualization, Supervision, Funding Acquisition, Writing-Original draft preparation, Writing-Reviewing and Editing.

## Competing interests

The authors declare no competing or financial interests.

## Notes

### Competing Interest Statement

The authors have declared no competing interest.

## References

(1) Andrew, D. J.; Ewald, A. J. Morphogenesis of Epithelial Tubes: Insights into Tube Formation, Elongation, and Elaboration. Dev. Biol. 2010, 341 (1), 34–55. 10.1016/j.ydbio.2009.09.024.

(2) Friedl, P.; Gilmour, D. Collective Cell Migration in Morphogenesis, Regeneration and Cancer. Nat. Rev. Mol. Cell Biol. 2009, 10 (7), 445–457. 10.1038/nrm2720.

(3) Ridley, A. J. Rho GTPase Signalling in Cell Migration. Curr. Opin. Cell Biol. 2015, 36, 103–112. 10.1016/j.ceb.2015.08.005.

(4) Lange, J. R.; Fabry, B. Cell and Tissue Mechanics in Cell Migration. Exp. Cell Res. 2013, 319 (16), 2418–2423. 10.1016/j.yexcr.2013.04.023.

(5) Song, B.; Guo, W.; He, Y.; Yao, X.; Sun, J.; Wang, S. Targeting Immune Cell Migration as Therapy for Inflammatory Disease: A Review. Front. Immunol. 2025, 16, 1650760. 10.3389/fimmu.2025.1650760.

(6) Trepat, X.; Chen, Z.; Jacobson, K. Cell Migration. Compr. Physiol. 2012, 2 (4), 2369–2392. 10.1002/cphy.c110012.

(7) Garcin, C.; Straube, A. Microtubules in Cell Migration. Essays Biochem. 2019, 63 (5), 509–520. 10.1042/EBC20190016.

(8) Seetharaman, S.; Etienne-Manneville, S. Cytoskeletal Crosstalk in Cell Migration. Trends Cell Biol. 2020, 30 (9), 720–735. 10.1016/j.tcb.2020.06.004.

(9) Zanotelli, M. R.; Zhang, J.; Reinhart-King, C. A. Mechanoresponsive Metabolism in Cancer Cell Migration and Metastasis. Cell Metab. 2021, 33 (7), 1307–1321. 10.1016/j.cmet.2021.04.002.

(10) Srinivasan, D.; Balakrishnan, R.; Chauhan, A.; Kumar, J.; Girija, D. M.; Shrestha, R.; Shrestha, R.; Subbarayan, R. Epithelial-Mesenchymal Transition in Cancer: Insights Into Therapeutic Targets and Clinical Implications. MedComm 2025, 6 (9), e70333. 10.1002/mco2.70333.

(11) Yamazaki, D.; Kurisu, S.; Takenawa, T. Regulation of Cancer Cell Motility through Actin Reorganization. Cancer Sci. 2005, 96 (7), 379–386. 10.1111/j.1349-7006.2005.00062.x.

(12) Mondal, C.; Di Martino, J. S.; Bravo-Cordero, J. J. Actin Dynamics during Tumor Cell Dissemination. Int. Rev. Cell Mol. Biol. 2021, 360, 65–98. 10.1016/bs.ircmb.2020.09.004.

(13) Nagano, M.; Hoshino, D.; Koshikawa, N.; Akizawa, T.; Seiki, M. Turnover of Focal Adhesions and Cancer Cell Migration. Int. J. Cell Biol. 2012, 2012, 310616. 10.1155/2012/310616.

(14) Samuel, M. S.; Lopez, J. I.; McGhee, E. J.; Croft, D. R.; Strachan, D.; Timpson, P.; Munro, J.; Schröder, E.; Zhou, J.; Brunton, V. G.; Barker, N.; Clevers, H.; Sansom, O. J.; Anderson, K. I.; Weaver, V. M.; Olson, M. F. Actomyosin-Mediated Cellular Tension Drives Increased Tissue Stiffness and β-Catenin Activation to Induce Epidermal Hyperplasia and Tumor Growth. Cancer Cell 2011, 19 (6), 776–791. 10.1016/j.ccr.2011.05.008.

(15) Nagano, M.; Hoshino, D.; Koshikawa, N.; Akizawa, T.; Seiki, M. Turnover of Focal Adhesions and Cancer Cell Migration. Int. J. Cell Biol. 2012, 2012, 1–10. 10.1155/2012/310616.

(16) Gandalovičová, A.; Rosel, D.; Fernandes, M.; Veselý, P.; Heneberg, P.; Čermák, V.; Petruželka, L.; Kumar, S.; Sanz-Moreno, V.; Brábek, J. Migrastatics—Anti-Metastatic and Anti-Invasion Drugs: Promises and Challenges. Trends Cancer 2017, 3 (6), 391–406. 10.1016/j.trecan.2017.04.008.

(17) Yayan, J.; Franke, K.-J.; Berger, M.; Windisch, W.; Rasche, K. Adhesion, Metastasis, and Inhibition of Cancer Cells: A Comprehensive Review. Mol. Biol. Rep. 2024, 51 (1), 165. 10.1007/s11033-023-08920-5.

(18) Al-Bari, M. A. A. Chloroquine Analogues in Drug Discovery: New Directions of Uses, Mechanisms of Actions and Toxic Manifestations from Malaria to Multifarious Diseases. J. Antimicrob. Chemother. 2015, 70 (6), 1608–1621. 10.1093/jac/dkv018.

(19) Haładyj, E.; Sikora, M.; Felis-Giemza, A.; Olesińska, M. Antimalarials - Are They Effective and Safe in Rheumatic Diseases? Reumatologia 2018, 56 (3), 164–173. 10.5114/reum.2018.76904.

(20) Plantone, D.; Koudriavtseva, T. Current and Future Use of Chloroquine and Hydroxychloroquine in Infectious, Immune, Neoplastic, and Neurological Diseases: A Mini-Review. Clin. Drug Investig. 2018, 38 (8), 653–671. 10.1007/s40261-018-0656-y.

(21) Abdel-Aziz, A. K.; Saadeldin, M. K.; Salem, A. H.; Ibrahim, S. A.; Shouman, S.; Abdel-Naim, A. B.; Orecchia, R. A Critical Review of Chloroquine and Hydroxychloroquine as Potential Adjuvant Agents for Treating People with Cancer. Future Pharmacol. 2022, 2 (4), 431–443. 10.3390/futurepharmacol2040028.

(22) Montanari, F.; Lu, M.; Marcus, S.; Saran, A.; Malankar, A.; Mazumder, A. A Phase II Trial of Chloroquine in Combination with Bortezomib and Cyclophosphamide in Patients with Relapsed and Refractory Multiple Myeloma. Blood 2014, 124 (21), 5775–5775. 10.1182/blood.V124.21.5775.5775.

(23) Verbaanderd, C.; Maes, H.; Schaaf, M. B.; Sukhatme, V. P.; Pantziarka, P.; Sukhatme, V.; Agostinis, P.; Bouche, G. Repurposing Drugs in Oncology (ReDO)-Chloroquine and Hydroxychloroquine as Anti-Cancer Agents. Ecancermedicalscience 2017, 11, 781. 10.3332/ecancer.2017.781.

(24) Sarkar, S.; Bisoi, A.; Singh, P. C. Spectroscopic and Molecular Dynamics Aspect of Antimalarial Drug Hydroxychloroquine Binding with Human Telomeric G-Quadruplex. J. Phys. Chem. B 2022, 126 (28), 5241–5249. 10.1021/acs.jpcb.2c03267.

(25) Sarkar, S.; Chatterjee, A.; Paul, S.; Bisoi, A.; Sen, P.; Chandra Singh, P. G-Quadruplex-Specific Action of Chloroquine-Based Immunomodulator Drugs to Inhibit the Cancer Progression. J. Biol. Chem. 2025, 301 (11), 110753. 10.1016/j.jbc.2025.110753.

(26) Kauanova, S.; Urazbayev, A.; Vorobjev, I. The Frequent Sampling of Wound Scratch Assay Reveals the “Opportunity” Window for Quantitative Evaluation of Cell Motility-Impeding Drugs. Front. Cell Dev. Biol. 2021, 9, 640972. 10.3389/fcell.2021.640972.

(27) Pollard, T. D.; Borisy, G. G. Cellular Motility Driven by Assembly and Disassembly of Actin Filaments. Cell 2003, 112 (4), 453–465. 10.1016/S0092-8674(03)00120-X.

(28) Fuseler, J. W.; Millette, C. F.; Davis, J. M.; Carver, W. Fractal and Image Analysis of Morphological Changes in the Actin Cytoskeleton of Neonatal Cardiac Fibroblasts in Response to Mechanical Stretch. Microsc. Microanal. Off. J. Microsc. Soc. Am. Microbeam Anal. Soc. Microsc. Soc. Can. 2007, 13 (2), 133–143. 10.1017/S1431927607070225.

(29) Ray, D.; Sinha, D. K. Dynamic Crosstalk between Cytoskeletal Filaments Regulates Dorsoventral Cytoplasmic Mechanics. J. Cell Sci. 2025, 138 (4), JCS263464. 10.1242/jcs.263464.

(30) Coelho-Rato, L. S.; Parvanian, S.; Modi, M. K.; Eriksson, J. E. Vimentin at the Core of Wound Healing. Trends Cell Biol. 2024, 34 (3), 239–254. 10.1016/j.tcb.2023.08.004.

(31) Murashko, A. V.; Frolova, A. A.; Akovantseva, A. A.; Kotova, S. L.; Timashev, P. S.; Efremov, Y. M. The Cell Softening as a Universal Indicator of Cell Damage during Cytotoxic Effects. Biochim. Biophys. Acta BBA - Gen. Subj. 2023, 1867 (6), 130348. 10.1016/j.bbagen.2023.130348.

(32) Mason, T. G. Estimating the Viscoelastic Moduli of Complex Fluids Using the Generalized Stokes-Einstein Equation. Rheol. Acta 2000, 39 (4), 371–378. 10.1007/s003970000094.

(33) Roopnarine, O.; Thomas, D. D. Mechanistic Analysis of Actin-Binding Compounds That Affect the Kinetics of Cardiac Myosin-Actin Interaction. J. Biol. Chem. 2021, 296, 100471. 10.1016/j.jbc.2021.100471.

(34) Strzelecka-Golaszewska, H.; Venyaminov, S. Yu.; Zmorzyǹski, S.; Mossakowska, M. Effects of Various Amino Acid Replacements on the Conformational Stability of G-actin. Eur. J. Biochem. 1985, 147 (2), 331–342. 10.1111/j.1432-1033.1985.tb08754.x.

(35) Doolittle, L. K.; Rosen, M. K.; Padrick, S. B. Measurement and Analysis of in Vitro Actin Polymerization. Methods Mol. Biol. Clifton NJ 2013, 1046, 273–293. 10.1007/978-1-62703-538-5_16.

(36) Zelnak, A. B. Clinical Pharmacology and Use of Microtubule-Targeting Agents in Cancer Therapy. Methods Mol. Med. 2007, 137, 209–234. 10.1007/978-1-59745-442-1_15.

(37) Sarkar, S.; Bisoi, A.; Singh, P. C. Antimalarial Drugs Induce the Selective Folding of Human Telomeric G-Quadruplex in a Cancer-Mimicking Microenvironment. J. Phys. Chem. B 2023, 127 (30), 6648–6655. 10.1021/acs.jpcb.3c03042.

(38) Edelstein, A. D.; Tsuchida, M. A.; Amodaj, N.; Pinkard, H.; Vale, R. D.; Stuurman, N. Advanced Methods of Microscope Control Using μManager Software. J. Biol. Methods 2014, 1 (2), e10. 10.14440/jbm.2014.36.

(39) Karperien, A. FracLac for ImageJ; NIH, 2013. 10.13140/2.1.4775.8402.

(40) Tseng, Y.; Kole, T. P.; Wirtz, D. Micromechanical Mapping of Live Cells by Multiple-Particle-Tracking Microrheology. Biophys. J. 2002, 83 (6), 3162–3176. 10.1016/S0006-3495(02)75319-8.

(41) Wirtz, D. Particle-Tracking Microrheology of Living Cells: Principles and Applications. Annu. Rev. Biophys. 2009, 38, 301–326. 10.1146/annurev.biophys.050708.133724. 10.1146/annurev.biophys.050708.133724.

(42) Sbalzarini, I. F.; Koumoutsakos, P. Feature Point Tracking and Trajectory Analysis for Video Imaging in Cell Biology. J. Struct. Biol. 2005, 151 (2), 182–195. 10.1016/j.jsb.2005.06.002.

(43) Bolte, S.; Cordelières, F. P. A Guided Tour into Subcellular Colocalization Analysis in Light Microscopy. J. Microsc. 2006, 224 (Pt 3), 213–232. 10.1111/j.1365-2818.2006.01706.x.

