## Supplementary information for "Hydroxychloroquine alters the cytoskeleton to impair cell migration"

**List of supplementary materials included:**

- **Supplementary Figure S1**
- **Supplementary Figure S2**
- **Supplementary Figure S3**
- **Supplementary Figure S4**
- **Video 1***
- **Video 2***

***Video files are uploaded separately. Legends of the videos are included at the end of this document.**

### Supplementary figures

#### Supplementary Figure S1

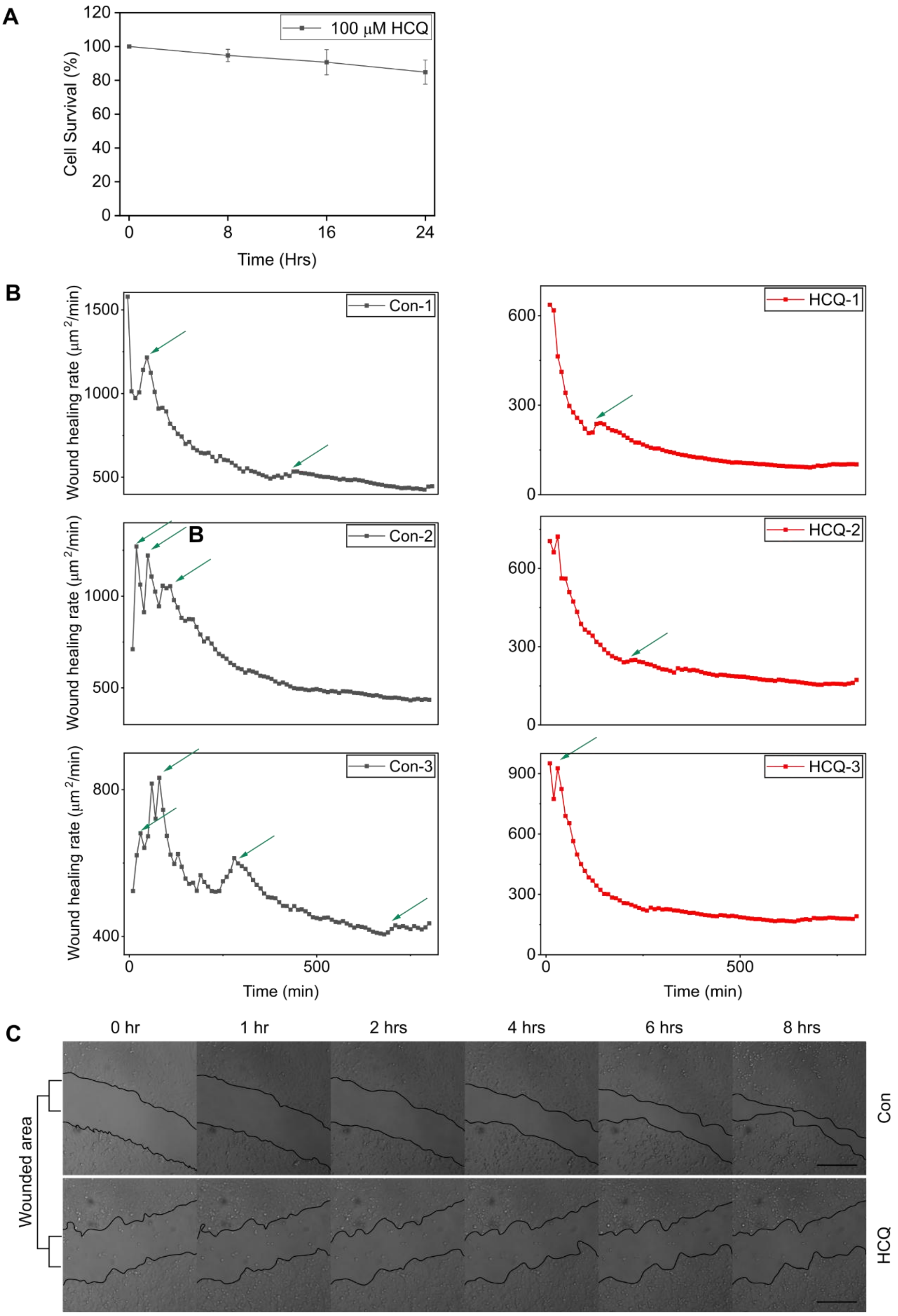

**Figure S1.**

(A) MTT assay showing the effect of 100 µM hydroxychloroquine (HCQ) on HeLa cell viability following treatment for 0, 8, 16, and 24 h. Cell survival was quantified relative to untreated control cells at 0 h, normalized to 100%.

(B) The graph displays the measured wound healing rates from individual, independent experimental replicates (sets) for both Control and HCQ-treated cell populations. Green arrows indicate intermittent spikes in the instantaneous wound healing rate observed over the measurement period.

(C) Wound healing assay comparing control and HCQ-treated MDA-MB-231 cells demonstrates markedly reduced migration with HCQ exposure. Representative phase-contrast images at 0, 1, 2, 4, 6 and 8 hrs show noticeably slower wound closure in HCQ-treated cells (scale bar, 200 µm).

**Supplementary Figure S2**

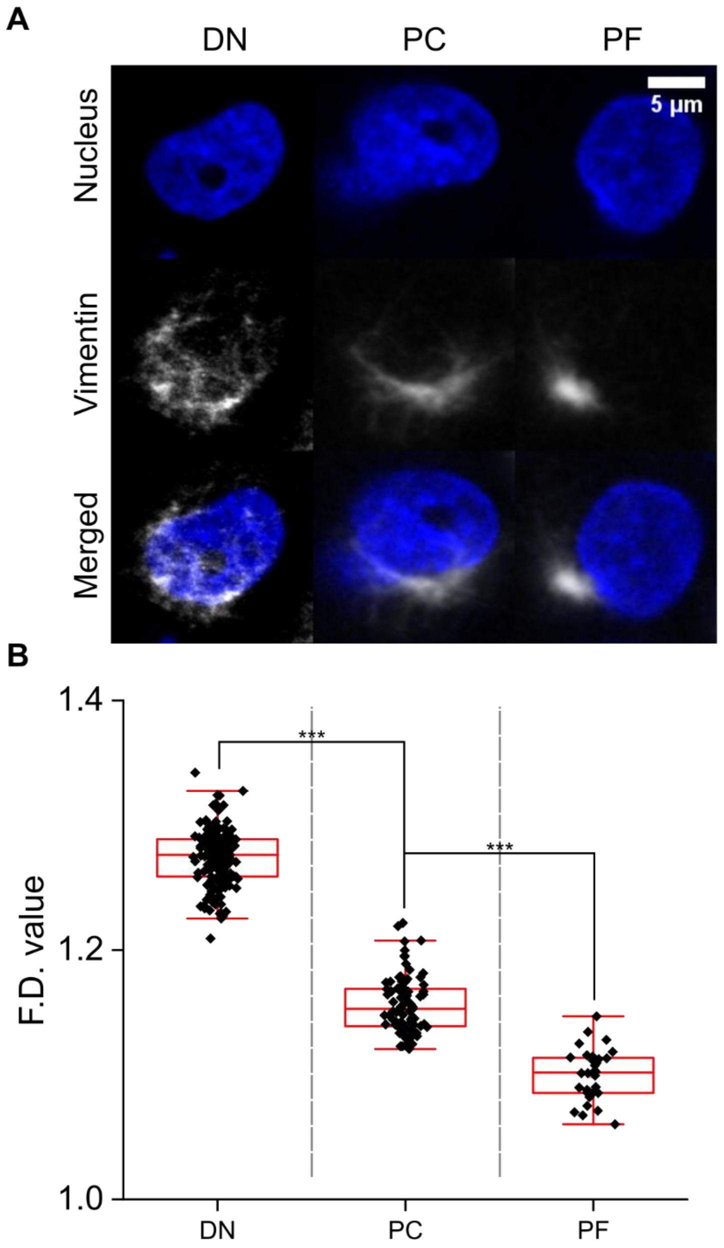


**Figure S2:** (A) Representative images showing vimentin localization in three different nuclear regions: dispersed network (DN), perinuclear cage (PC), and perinuclear foci (PF). The top row displays nuclear staining (blue), the middle row shows vimentin immunofluorescence (white), and the bottom row shows the merged images of both (blue and white). Scale bar, 5 µm (B) Boxplot showing the fractal dimension (F.D. value) of vimentin in the DN, PC, and PF regions. Higher F.D. values correspond to a denser vimentin network ($N_{cells}$>20, ***p<0.001; unpaired t-test)

**Supplementary Figure S3**

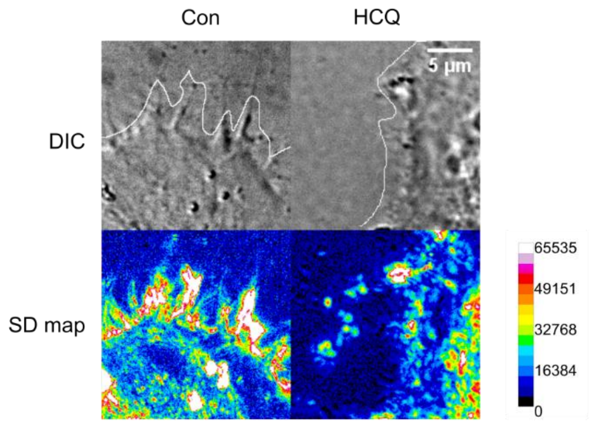


**Figure S3:** Standard deviation (SD) map of the lamellipodial regions of single control and HCQ-treated HeLa cells. Scale bar, 5 µm.

#### Supplementary Figure S4

(a)

| | $\alpha_1$ | $\tau_1$ | $\alpha_1$ | $\tau_2$ | $\tau_{avg}$ | $\chi^2$ |
|---|---|---|---|---|---|---|
| Actin | 0.42 | 1.61 | 0.58 | 4.94 | 3.54 | 1.18 |
| +HCQ | | | | | | |
| 2 µM | 0.61 | 1.18 | 0.39 | 3.26 | 1.99 | 1.12 |
| 5 µM | 0.51 | 1.02 | 0.49 | 2.20 | 1.59 | 1.07 |
| 10 µM | 0.17 | 0.42 | 0.83 | 1.65 | 1.44 | 1.11 |
| 20 µM | 0.12 | 0.37 | 0.88 | 1.62 | 1.47 | 1.03 |
| 30 µM | 0.13 | 0.36 | 0.87 | 1.63 | 1.46 | 1.09 |
| 50 µM | 0.13 | 0.16 | 0.87 | 1.60 | 1.41 | 1.07 |

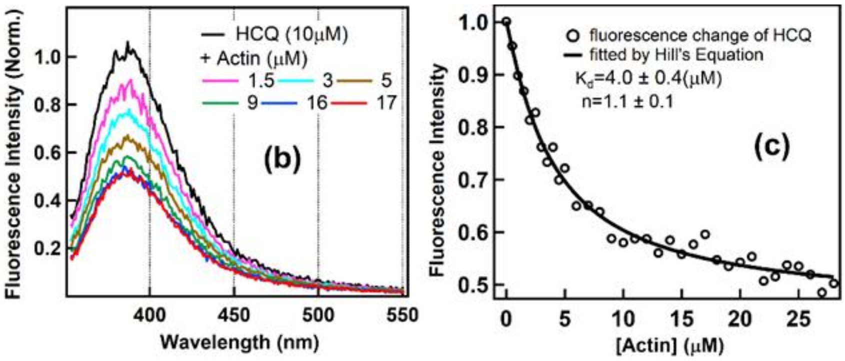


**Figure S4**: (a) The amplitude and the value of the lifetime components of actin (10 µM) in the absence and presence of HCQ. (b) The fluorescence spectra of HCQ (10 µM) in the absence and presence of actin obtained on the excitation of HCQ only at 340 nm. (c) The plot of the change in the emission maxima of HCQ (~380 nm) with the addition of the actin. The $K_d$ value of the binding of actin with HCQ was obtained using the Hill's equation.

**Video 1:** DIC time-lapse video of control cells undergoing a wound healing assay. The time interval between each frame is 10 minutes. Scale bar: 200 µm.

**Video 2:** Time-lapse video of nuclei in control cells stained with Hoechst 33342. The time interval between successive frames is 2 minutes. Scale bar: 200 µm.
